# Emergence of opposite neurons in a decentralized firing-rate model of multisensory integration

**DOI:** 10.1101/845743

**Authors:** Xueyan Niu, Ho Yin Chau, Tai Sing Lee, Wen-Hao Zhang

## Abstract

Multisensory integration areas such as dorsal medial superior temporal (MSTd) and ventral intraparietal (VIP) areas in macaques combine visual and vestibular cues to produce better estimates of self-motion. Congruent and opposite neurons, two types of neurons found in these areas, prefer congruent inputs and opposite inputs from the two modalities, respectively. A recently proposed computational model of congruent and opposite neurons reproduces their tuning properties and shows that congruent neurons optimally integrate information while opposite neurons compute disparity information. However, the connections in the network are fixed rather than learned, and in fact the connections of opposite neurons, as we will show, cannot arise from Hebbian learning rules. We therefore propose a new model of multisensory integration in which congruent neurons and opposite neurons emerge through Hebbian and anti-Hebbian learning rules, and show that these neurons exhibit experimentally observed tuning properties.

## Introduction

Multisensory integration is the task of combining information about an external stimulus gathered from different sensory modalities in order to improve perception. For example, information about heading direction may come from both visual inputs (optic flow) and vestibular inputs (self-motion), and it is therefore useful to integrate this information together to produce a better estimate of heading direction. Humans integrate multisensory information in a near-optimal way according to Bayes’ rule, and it is desirable to understand how this is performed by underlying neural circuits.

The multisensory neurons in visual and vestibular brain areas, such as dorsal medial superior temporal area (MSTd) and the ventral intraparietal (VIP) areas, can be divided into two categories according to their tuning properties. One type of neurons is called congruent neuron, as they prefer visual and vestibular cues of the same heading directions (Fig. 3C, tuning curve of congruent neurons). The other type is opposite neuron, which prefer visual and vestibular cues of opposite heading directions (Fig. 3D, tuning curve of opposite neurons) [1, 7, 8, 14, 18]. Congruent neurons have been proposed to be the neural basis of multisensory integration in monkeys, but the functional significance of opposite neurons is less clear [1]. It has been recently hypothesized that these neurons are involved in the decision of whether to integrate or segregate different sensory information based on the likelihood that these cues have a common cause [30, 31]. This serves as an important computational role as it does not make sense to integrate sensory information that have different causes. For example, if a person is wearing a virtual reality headset but sitting still, then visual and vestibular cues of heading direction would be inconsistent and the brain should not integrate the two cues.

Because of the potential significance of their computational function, it is desirable to build a model of multisensory integration with congruent and opposite neurons to achieve this function. One such model is the decentralized multisensory integration model proposed recently [30]. This model is able to account for the tuning properties of congruent and opposite neurons, and it further demonstrates that multisensory integration can be performed near-optimally in the model. However, the synaptic connections in the model are fixed rather than learned. A more serious problem, however, is that the design imposed on opposite neurons cannot arise from Hebbian learning rules in a natural way. In this study we propose an alternative neural circuit that can successfully learn congruent and opposite neurons with biologically realistic learning rules, and demonstrate that the learned neurons have tuning properties that agree with experiments as well as theoretical predictions from probabilistic inference.

## Results

### Hebbian learning fails to learn tunings opposite to statistics of natural scenes

We consider a neural circuit model with synaptic plasticity to learn congruent and opposite neurons in MSTd and VIP which receive both visual and vestibular stimuli. Congruent and opposite neurons are named after their tuning properties: congruent neurons prefer visual and vestibular stimuli under similar heading directions, while opposite neurons prefer visual and vestibular stimuli under opposite heading directions, i.e. heading directions differing by 180° (Fig. 3D). Previous network models (e.g., [30], [13]) propose that congruent and opposite tunings could emerge from combining excitatory inputs from two sensory modalities in a congruent or opposite manner respectively. For example, a congruent neuron preferring 0° visual motion would receive excitatory inputs at 0° from both sensory modalities, while an opposite neuron preferring 0° visual motion would receive excitatory visual inputs at 0° and excitatory vestibular inputs at 180°. These models could reproduce a wide range of neurophysiological observations on congruent and opposite neurons.

Although the connectivity scheme in these previous models is simple and intuitive, a serious problem occurs when we attempted to learn the opposite tunings with a Hebbian learning rule in a world where most visual and vestibular directions are consistent with each other. We simulated a population of excitatory neurons with Hebbian rule (Fig. 1A) that receive inputs with joint input statistics as shown in Fig. 1B, **which we believe is a proper assumption of the statistics of natural scenes**. After learning, all of the neurons developed congruent tunings to visual and vestibular stimuli, and no opposite neurons emerged in this network, as shown in Fig. 1C, opposed to approximately the same number of congruent and opposite neurons found in previous experiments [14] (Fig. 1D). This is because the Hebbian rule learns to form associations between visual and vestibular cues that are most correlated. In a world with consistent visual and vestibular directions, the two cues are highly correlated, and therefore neurons form congruent tunings but not opposite tunings.

**Fig 1.**
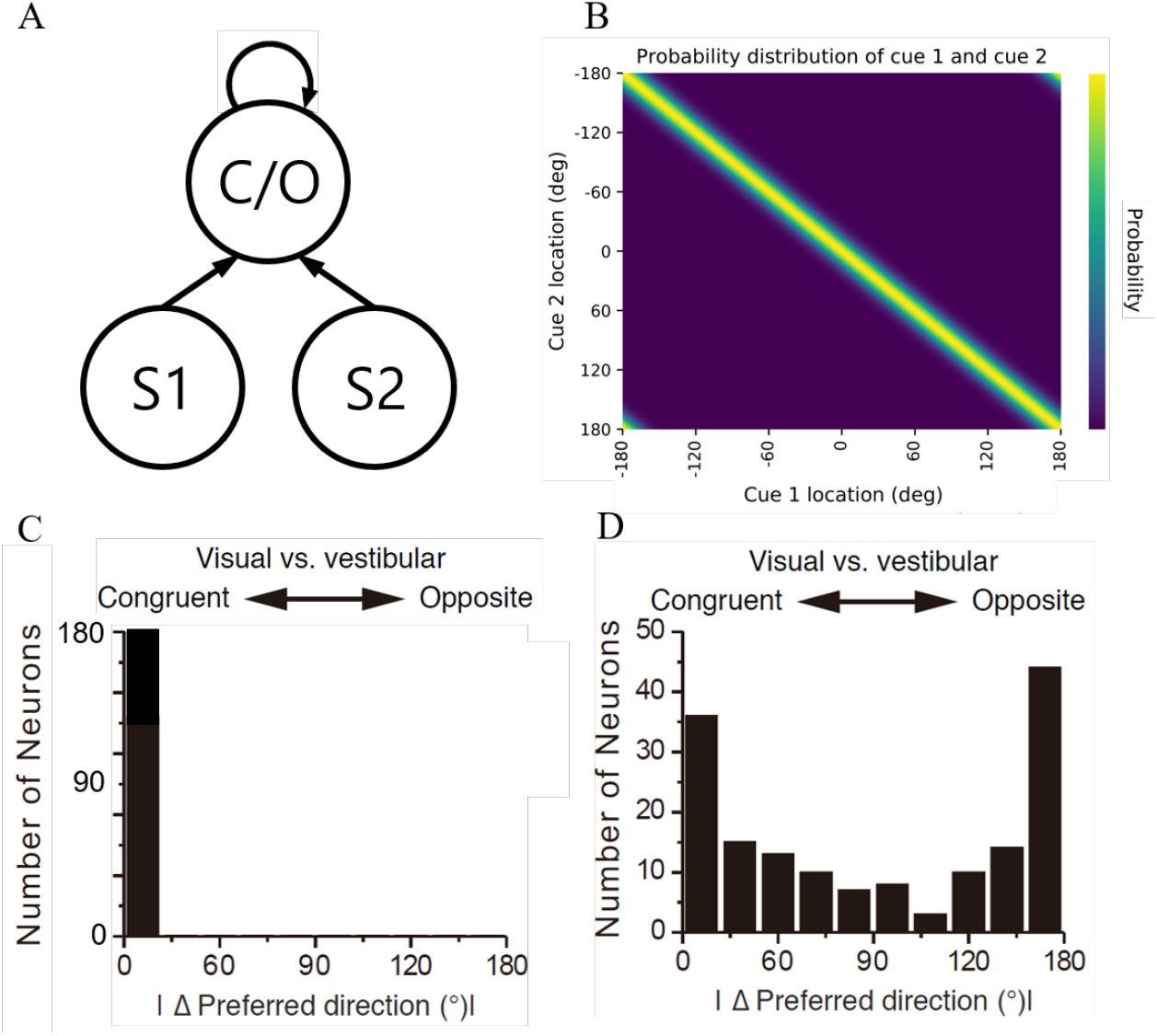
Hebbian learning alone cannot explain emergence of opposite neurons. A) Network model architecture used in this simulation, where congruent and opposite neurons receive direct, excitatory connections from S1 and S2 inputs. B) Correlation of S1 and S2 inputs used in our simulation. C) Distribution of congruent and opposite neurons within the learned network with Hebbian learning only. D) Distribution of congruent and opposite neurons in the macaque MSTd, from [14].

Is it possible that the failure to learn opposite neurons comes from a wrong assumption about the joint distribution of visual and vestibular direction in our study? We performed a control experiment in which most visual and vestibular directions are opposite, and we found the opposite neurons emerge while the congruent neurons disappear. In other words, the simple Hebbian mechanism still fails to learn excitatory congruent and opposite tunings simultaneously. Moreover, we believe the visual and vestibular directions the brain receives are mostly similar instead of opposite with each other, although no work so far has studied the joint statistics of visual and vestibular directions received by the brain. This is because the vestibular direction represents our self-motion direction and the visual direction is a mixture of self-motion and the direction of other moving objects. Although the moving objects contribute to certain discordance between visual and vestibular directions, it is very unlikely that most of the objects move in the same direction as our self-motion and faster than us (hence contradict to the relative motion of static background). Therefore it is highly unlikely that the failure of learning opposite neurons results from a wrong assumption about the joint distribution of input directions. This motivated us to consider a new network framework with biologically plausible synaptic plasticity rule from which the congruent and opposite neurons can emerge simultaneously from the input distribution as in Fig. 1B.

### A biologically learnable decentralized architecture of congruent and opposite neurons

As above analysis indicates, the Hebbian rule is not able to learn the opposite connections involved in previously proposed models of congruent and opposite neurons. Here we propose an alternative network structure in which the opposite tunings can be learned without the need for opposite connections, as depicted in Fig. 2A. Opposite neurons in this model have congruent instead of opposite connections, in the sense that each opposite neuron may, for example, receive inhibitory input from congruent neurons with preferred motion direction *θ* and excitatory visual input at *θ* as well, whereas in previous models the opposite neuron may receive excitatory visual input at *θ* and excitatory vestibular input at *θ* + 180°, which cannot be spontaneously learned by Hebbian rules.

**Fig 2.**
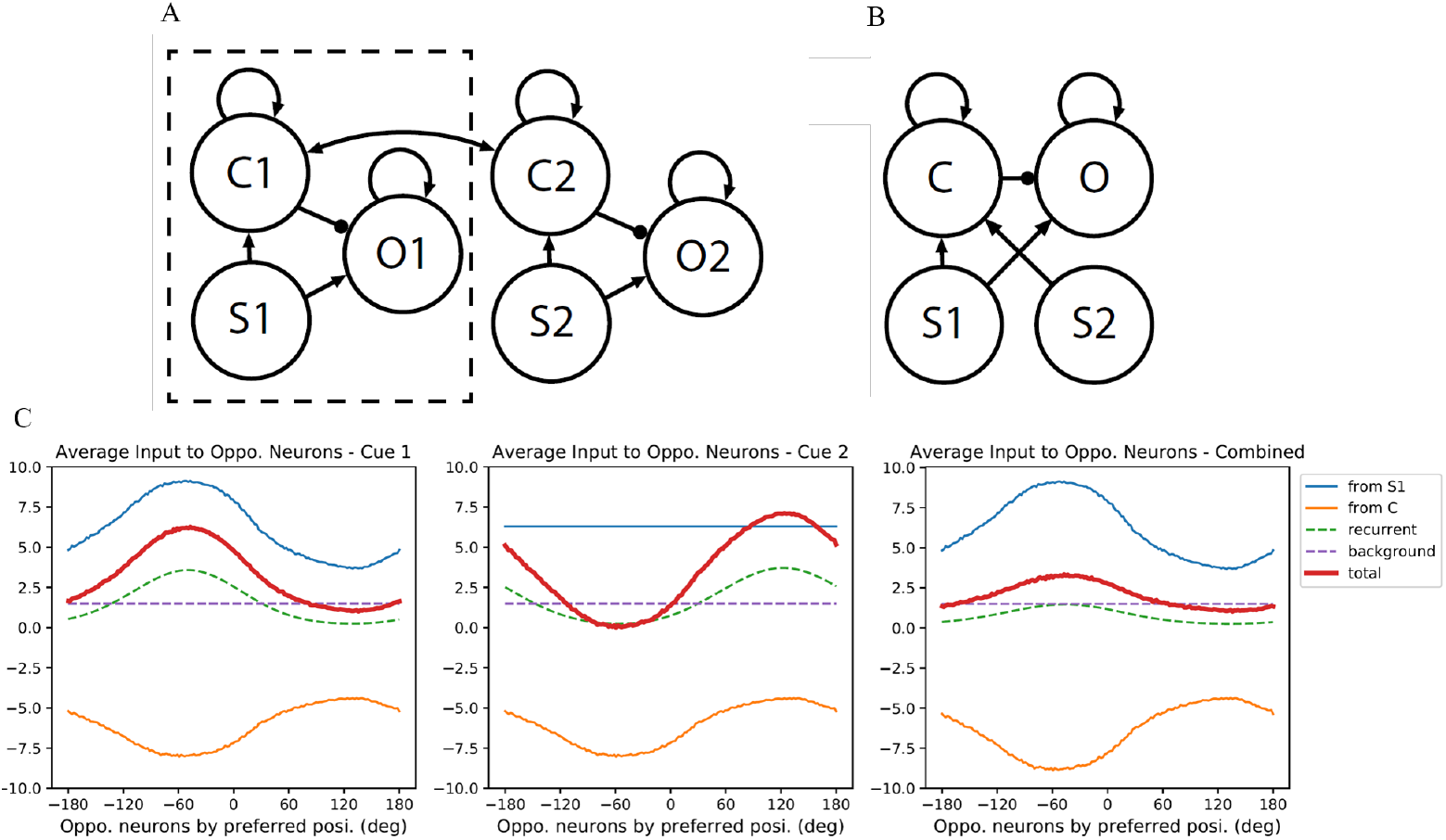
Network architecture and mechanism of opposite tuning. A) Our model architecture. Each sensory modality has its own uni-sensory neurons (S1, S2) as well as congruent and opposite neurons (C1, C2, O1, O2), with excitatory reciprocal connections bridging the two modules. Feedforward connections and recurrent connections are shown in the diagram, but divisive normalization is not shown. Arrow indicates excitatory connection, while a dot indicates inhibitory connection. Each group of neurons (S1, S2, C1, C2, O1, O2) are assumed to lie in a 1D ring formation, with their preferred direction ranging from [−*π*, *π*). B) A simplified model we tried first to validate our inhibitory connection proposal, which is equivalent to the boxed component in A). In this model, the excitatory connections were pre-set and we trained the inhibitory connection only. C) Based on the simplified model, an illustration on the input of opposite neurons in 3 conditions: only cue 1 present, only cue 2 present, both cue 1 and cue 2 present.

#### Opposite tuning is mediated by inhibition

How can opposite tuning be achieved without opposite connections? We show that opposite tuning emerges from the inhibition from congruent neurons to opposite neurons, conditioned on two simple and biologically plausible assumptions. The first assumption is that congruent and opposite neurons are broadly tuned to the heading direction, meaning the neurons are widely connected with each other on the ring, which is consistent with broad congruent and opposite tuning observed in experiments [14]. The second assumption is the opposite neurons receive a homogeneous background input larger than the peak inhibitory input from congruent neurons, in order to avoid the situation where all opposite neurons become silent after rectification.

Fig. 2A demonstrates the model architecture we propose. To separate the role of inhibition from congruent neurons to opposite neurons, we first discuss a simplified version of this model by discarding the second module (Fig. 2B) and examining the input to a population of opposite neurons **ordered by their preferred S1 direction** under 3 conditions: only cue 1 is presented (−60°), only cue 2 is presented (−60°) and both cues are presented (both are −60°). Specifically, when only cue 2 is presented, the middle figure of Fig. 2C shows that this input from S2 results in a broad inhibition of opposite neurons centered at −60° (yellow curve) due to excitation of congruent neurons at −60°. The background input (purple) and S1 excitation (uniform in all directions, blue, will be explained in section **Single neuron response**) balances out the large inhibition and causes total input to opposite neurons to be centered at 120° instead (red). As such, opposite neurons are tuned oppositely to S1 and S2 inputs.

Note that the proposed mechanism of inhibitory synapses from congruent neurons to opposite neurons is not inconsistent with experimental findings. Experiments only revealed that the opposite neurons exhibit facilitatory responses when inputs from two sensory modalities having opposite directions [18], however, which doesn’t necessarily mean the facilitatory responses are mediated by excitatory synaptic connections.

#### The inhibitory connections from congruent to opposite neurons can be learnt by anti-Hebbian rule

The only remaining question is how the congruent, inhibitory connections from congruent neurons to opposite neurons can be learned. We propose that these connections follow the anti-Hebbian rule (see Eq. 7), where correlated activity causes a reduction in connection weight. However, it can be more simply and intuitively understood as Hebbian rule on inhibitory interneurons, where correlated activity causes increase in the inhibitory synapses strength, effectively enhancing the inhibitory input. In fact, the learning rule we used for inhibitory connections has the same form as the learning rule for excitatory connections. As Hebbian learning results in congruent connections, we successfully learn congruent, inhibitory connections from congruent neurons to opposite neurons, with the two types of neurons function properly as in Fig. 3A-B.

**Fig 3.**
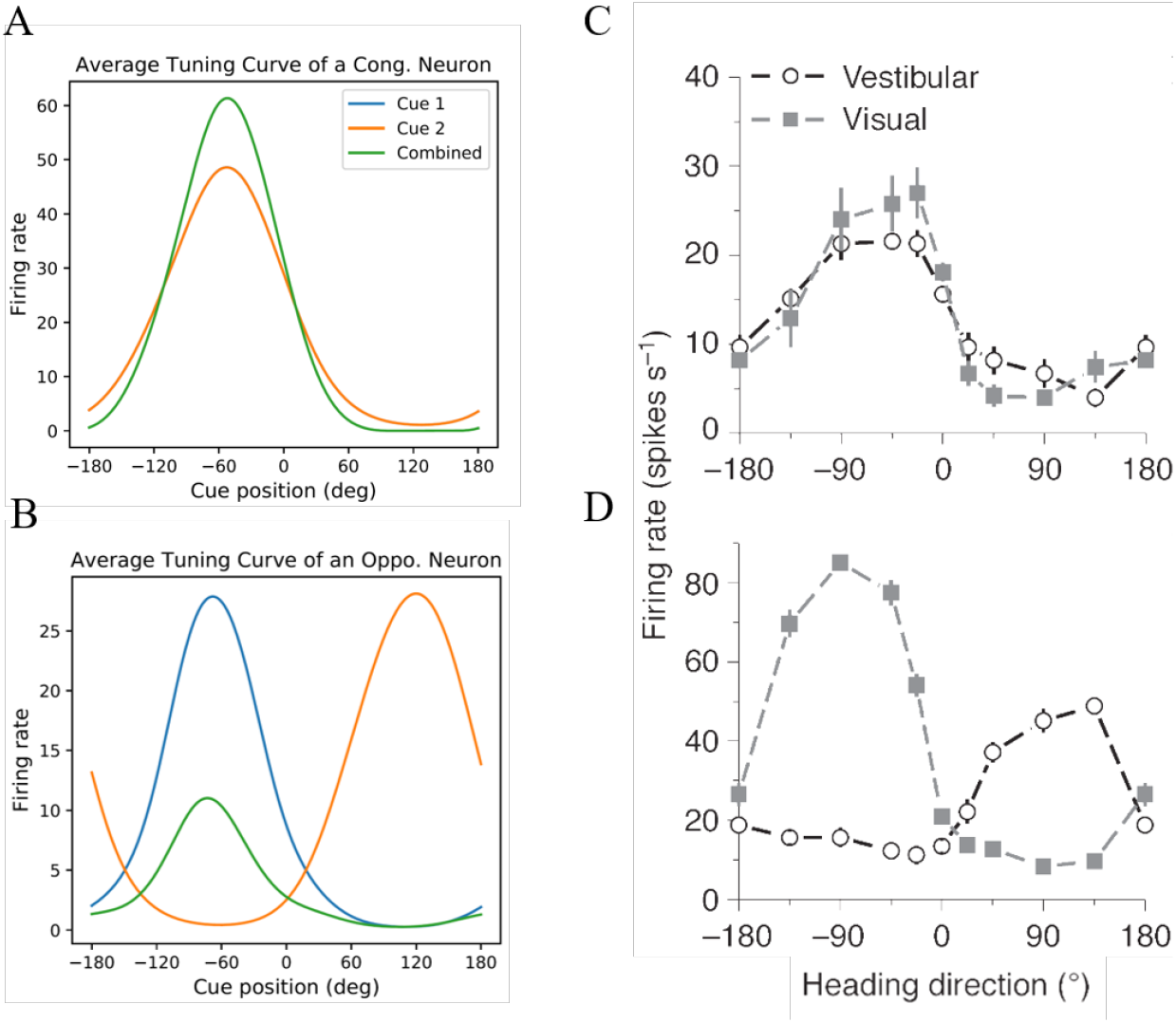
Tuning curves of one congruent neuron and one opposite neuron to unimodal and bimodal stimuli. A) Tuning curve of a congruent neuron when a) only cue 1 is presented, b) only cue 2 is presented, and c) both cues are presented at the same location. B) Similar, but for opposite neuron. C-D) Experimental data of tuning curves of MSTd neurons to vestibular and visual stimulus, reproduced from [1]. C and D show the tuning of a congruent neuron and an opposite neuron, respectively.

### Learned feedforward and reciprocal weights

Now we characterize the connections that are learned with our model dynamics and show that our network exhibits *self-organization*, where congruent and opposite neurons learn to be topographically organized with respect to their angles of tuning to S1 and S2 inputs.

Referring to Fig. 2A, we call the connections from *S_m_* to *C_m_*, from *S_m_* to *O_m_*, from *C_m_* to *O_m_* feedforward connections, with *M* ∈ {1, 2} representing the two modules. In particular, we call the connections from *S_m_* to *C_m_*, from *S_m_* to *O_m_* direct feedforward connections, as they indicate feedforward input directly from uni-sensory neurons. The connections within congruent neurons and opposte neurons are called recurrent connections, and the bi-directional connections between *C*1 and *C*2 are called reciprocal connections.

#### Feedforward connections to congruent and opposite neurons

We define the feedforward connections from S1/S2/C1/C2 neurons to some neuron *i* as the weight vector 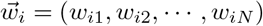, where *w_ij_* is the weight of a connection from a S1/S2/C1/C2 neuron *j* to neuron *i*. Effectively, 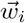 describes the receptive field of neuron *i*. The shape of feedforward connections 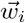 with respect to its indices *j* is found to be approximately proportional to a von-Mises distribution. This is shown in Fig. 4A-C. The assumption of Gaussian or von Mises shaped feedforward connections is usually assumed in multisensory integration models, and we show that the same shape can naturally come out in our learning model [9, 24, 25, 29, 30]. Since the model is completely symmetric over two modules, we will only show results of the first module.

**Fig 4.**
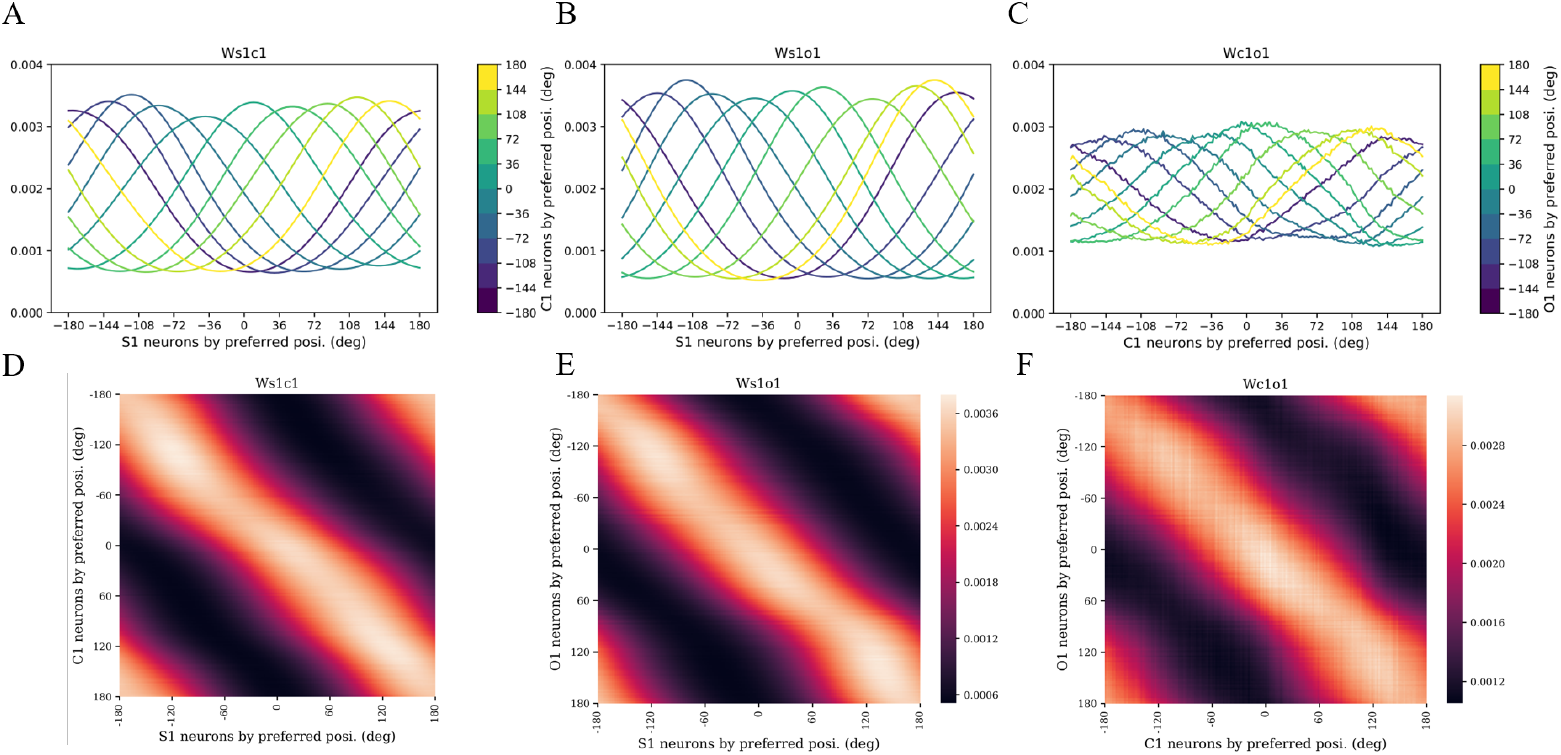
The learned feedforward connections and their topological order. A-C) Feedforward weights in module 1. Note that the weights in C) are the absolute value of the inhibitory connections learned by Anti-Hebbian rules. D-F) Topological order is naturally preserved due to recurrent input inside individual rings. The heatmap shows the strength of connection. In all cases, there is a bright diagonal, showing that neurons preferring the same direction in all pairs are strongly wired together.

Moreover, all pairs in Fig. 4A-C are congruently connected. For example, the yellow curve in Fig. 4C represents the opposite neuron in module 1 preferring 144°direction of cue 1 while it also receives the largest inhibition from the congruent neuron in module 1 preferring 144° direction of cue 1. The topological order is also well-preserved from uni-sensory neurons to congruent neurons and to opposite neurons, as shown in Fig. 4D-F. This organization naturally forms from the fact that the closer the neurons are in one ring, the stronger their recurrent connection is.

#### Reciprocal connections between two modules

It is not surprising that the reciprocal connections between two modules also have the shape of von-Mises distribution, as shown in Fig. 5A-B. Note that these connections are roughly five times smaller than feedforward connections, since the sum of recurrent input and reciprocal input for congruent neurons cannot exceed the critical value (see Materials and Methods) to prevent spontaneous activities of the two rings of congruent neurons. Therefore, the information from indirect cue (information from the other module) is much less than the direct cue (information from the self module), which will be demonstrated in the population response. C1 and C2 neurons are also topologically organized, as shown in Fig. 5C-D.

**Fig 5.**
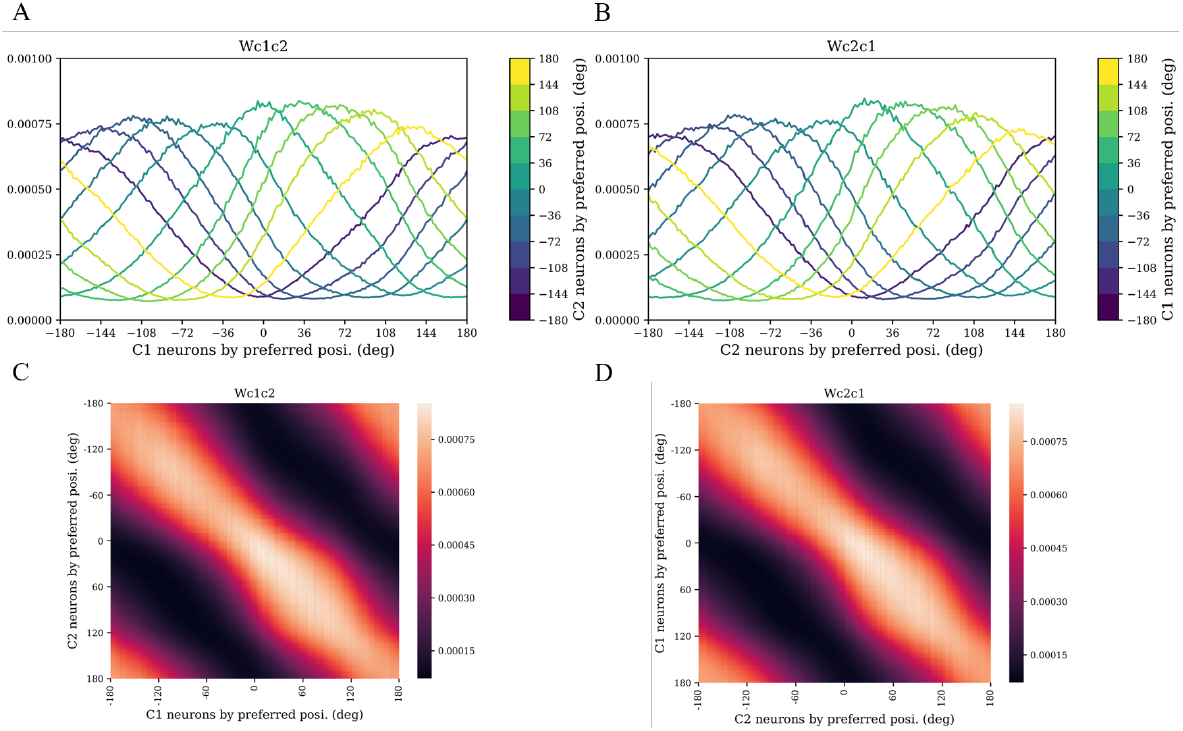
The learned reciprocal connections and their topological order. A) Reciprocal weights from module 1 to module 2. B) Reciprocal weights from module 2 to module 1. C-D) Topological order of congruent neurons bridging the two modules.

### Single neuron response

#### Tuning curves of congruent and opposite neurons

We mimicked neurophysiological experiments and obtained the tuning curves of congruent and opposite neurons by varying the input stimulus location and recording the response of the neurons. We applied both unimodal and bimodal stimuli, or the three conditions, and compared the resulting tuning curves. Here, a unimodal S1 stimulus (or “only cue 1 is presented”) means that the mean firing rate at S1 follows Eq. 4 with *R* = 1, while the mean firing rate at S2 follows the same equation with *R* = 0. Note that a unimodal S1 stimulus does not mean there is no input at S2, only that the input at S2 is a constant across all input directions. This is consistent with the observation that MT neurons (which would correspond to S1 or S2 in our model) appear to have a non-zero background input [4, 5]. Moreover, we note that for our model to work, a certain kind of homeostasis must be maintained: the total input from S1 and S2 has to remain relatively constant. This necessitates the use of a constant input at S2 even when we assume a unimodal S1 stimulus. A bimodal stimulus (or “both cues are presented”) means that both S1 and S2 neurons have the same mean firing rate with *z*_1_(*t*) = *z*_2_(*t*) in Eq. 4.

Fig. 6A-B shows the tuning curve of a neuron in C1 and a neuron in O1 that prefer a stimulus of 50° from modality 1. When only cue 1 is presented, the two neurons fire normally. When only cue 2 is presented, the neuron in C1 also prefers a stimulus of 50° from modality 2 but fires less, because the input from modality 2 is indirect and relatively weaker. When both cue 1 and cue 2 are presented, the tuning of the congruent neuron has a similar shape with stronger yet sub-additive response. For opposite neurons, tuning to cue 1 and cue 2 are separated by approximately 180°. When bimodal stimuli are presented, the response is flattened and highly sub-additive. The subadditivity of congruent and opposite neuron responses agree with experimental observations of MSTd neurons in macaques [18].

**Fig 6.**
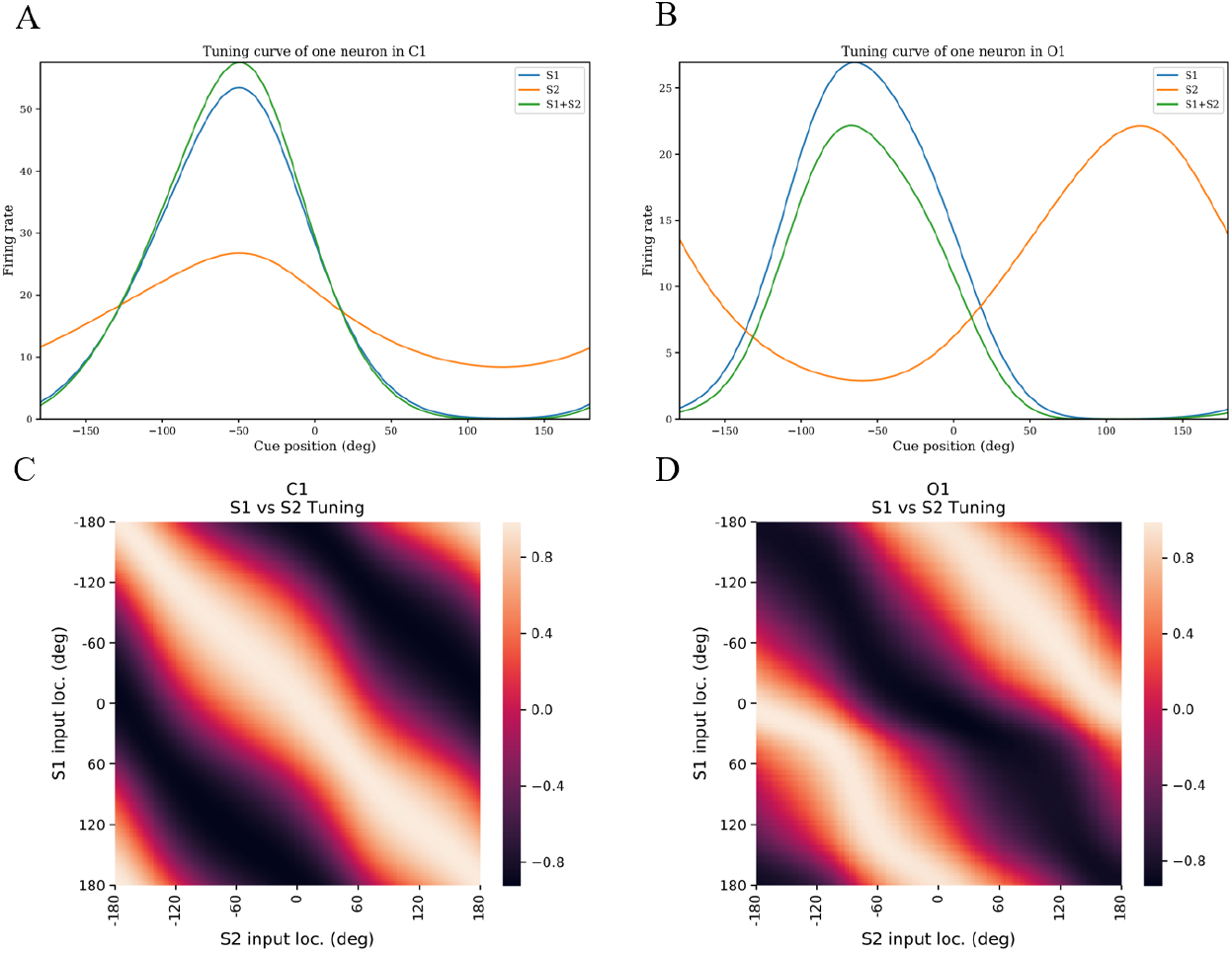
Tuning curves. A-B) Tuning curves for a congruent neuron and an opposite neuron preferring 50°cue from modality 1 under three conditions. C-D) Correlation of tuning curves of C1 neurons and O1 neurons towards unimodal S1 stimulus and unimodal S2 stimulus. For congruent neurons, we see a bright ridge along the main diagonal, meaning there is strong correlation of response to S1 and S2 inputs from the same location. For opposite neurons, we see a bright ridge along the diagonal shifted by 180°, meaning there is strong correlation of response to S1 and S2 inputs from opposite locations.

To show that congruent and opposite neurons have congruent and opposite tuning to S1 and S2 stimulus, we computed the correlation of their tuning curves towards unimodal S1 stimulus and unimodal S2 stimulus, as shown in Fig. 6C-D. For congruent neurons, responses to unimodal S1 and unimodal S2 stimuli are most strongly correlated when the inputs are at the same location, while for opposite neurons, responses are most strongly correlated when the inputs are separated by 180°.

#### Dependence of tuning on relative reliability of bimodal stimuli

Moreover, we show how the tuning of congruent and opposite neurons to both bimodal stimuli change as we decrease the reliability of one stimulus (Fig. 7). Physiological experiments have shown that as the reliability of one stimulus decrease, the neuron should be increasingly tuned to the other, more reliable stimulus [18]. This effect is also observed in our model.

**Fig 7.**
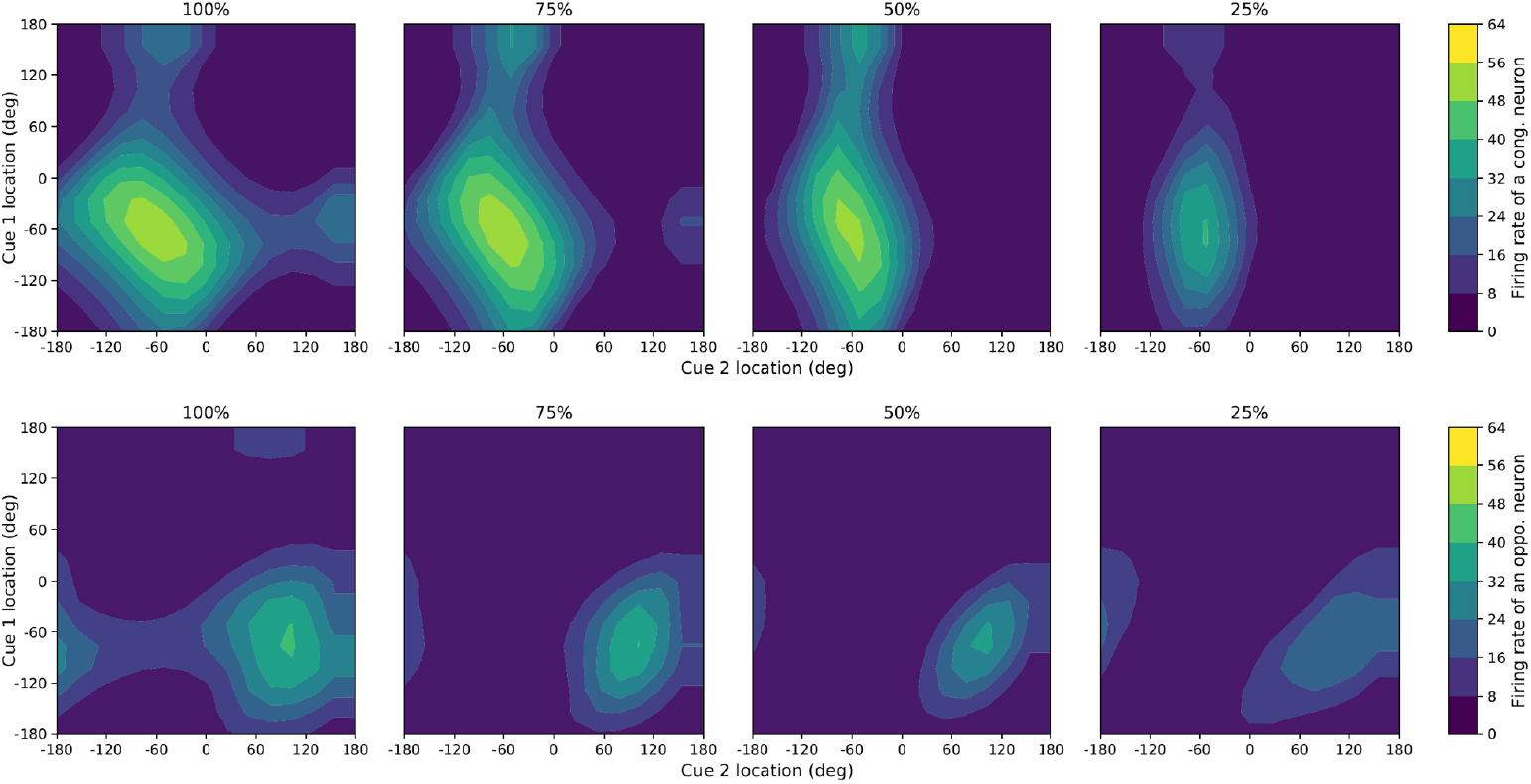
Dependence of tuning on relative reliability of bimodal stimuli. Change in tuning of a congruent and an opposite neuron to bimodal stimuli as S1 input reliability is decreased. As can be seen, decreasing S1 reliability shifts the tuning towards unimodal S2. Percentage indicates reliability of S1 input. Contour colors indicate firing rate.

### Collective response

#### Population response of congruent and opposite neurons

The population response of congruent neurons and opposite neurons when presented bimodal or unimodal stimuli is also qualitatively within expectation, as shown in Fig. 8, where the x-axis indicates neurons in the ring S1, S2, C1, C2, O1, O2, ordered by their preferred directions of their own modalities. Again, a unimodal S1 stimulus does not mean S2 has zero input but rather input with zero reliability, or a constant input. In figures below, S1 is centered at 0°, and S2 is centered at 60°.

**Fig 8.**
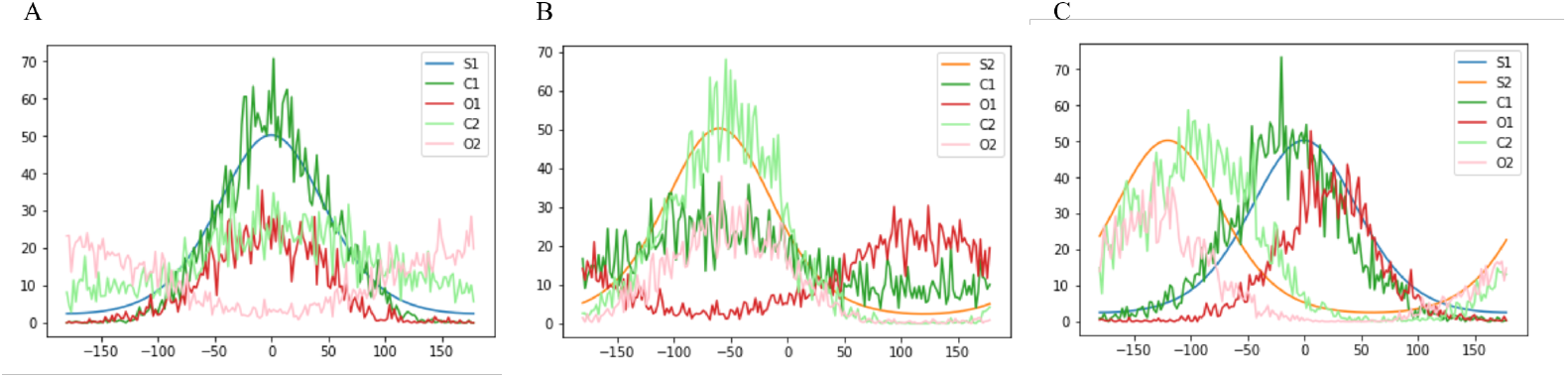
Population response of neurons in C1, O1, C2, O2. A) Only cue 1 is present. B) Only cue 2 is present. C) Both cues are present.

The peak of population response of congruent neurons is always in accordance with the center of unimodal stimulus S1 or S2. For opposite neurons, their peak of population response is the same as the center of input from their own modality, whereas opposite to that from the other modality.

When both cue 1 and cue 2 are presented, the congruent neurons seem to **integrate the two cues** and have the population response peak in between the center of S1 and S2, but biased by the input from their own modality. This is intuitive because the input from the other modality is indirect and relatively weaker. Interestingly, the opposite neurons seem to **reflect the disparity of the two cues**. The peak of the population response of O1 lies between S1 and the opposite of S2 (or S2 +180°), biased by S1, while the peak of O2 lies between S2 and the opposite of S1, biased by S2.

#### Comparing our decoding results with theoretical predictions

The peak of population response of C1, C2, O1, O2 in Fig. 8 is crucial in evaluating the performance of our model, as it reflects the brain’s **decoding**, or the neurons’ final decision of the heading direction after analyzing the two cues from different modalities. Since there is noise both in input and neural response, for each cue condition, we simulated the network 100 times and recorded the peaks of population response. The peaks turned out the form von-Mises distributions over the 100 trials, characterised by a mean and a concentration (essentially the inverse of variance, see **Materials and Methods**).

The paper by Zhang et al. [30] applies probabilistic inference and gives out the rules by which congruent neurons and opposite neurons should perform if the network achieves optimal integration, and we will use them to quantitatively assess our model.

Refer to [30], we use *y_l_*, *κ_l_* to denote the decoded mean and concentration of the population response, where *l* ∈ {*c*1,*c*2,*o*1,*o*2} separates congruent neurons and opposite neurons. In case of optimal integration, for congruent neurons, we should have

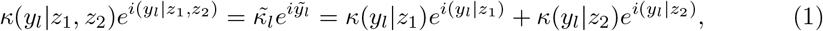

where *i* is the imaginary number, *z*_1_, *z*_2_ is the center of input at S1 and S2. The last term denotes the mean and concentration when the network is given unimodal stimulus, whose sum gives predictions to the mean and concentration when the network is given bimodal stimuli, namely 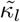 and 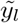, which we call **predicted decoding**. The first term is the **actual decoding** of the network, that is, the actual peak distribution when given bimodal stimuli. On the contrary, for opposite neurons, we would hope to have

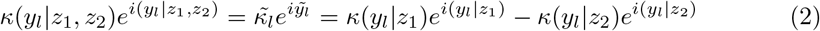

when *l* = *o*1, and

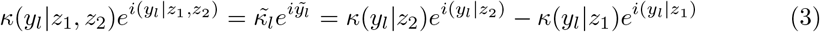

when *l* = *o*2, for optimal integration.

In the figure below, an S1 of 0°is given together with an S2 put at 30°intervals, ranging from [−*π*, *π*). The x-axis is the direction of S2. Our model can give promising predictions of the decoding of congruent neurons and opposite neurons, thus roughly achieved optimal integration.

**Fig 9.**
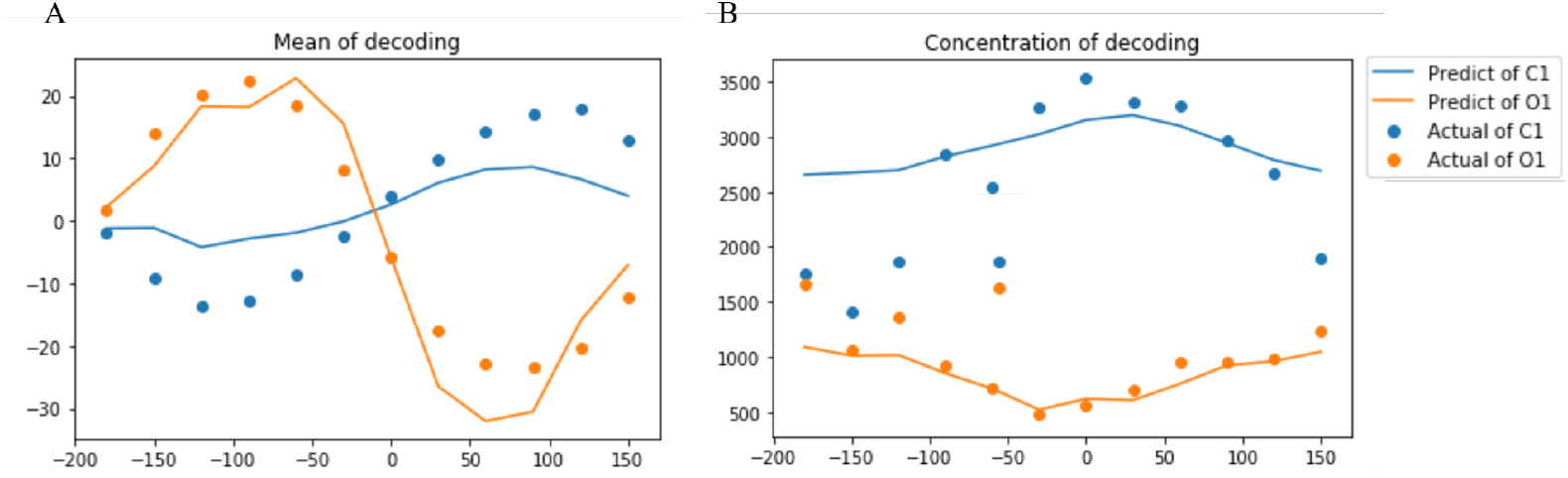
Decoding results. A) Actual and predicted mean of the peak distribution. B) Actual and predicted concentration of the peak distribution.

## Discussion

### Summary and relation to other works

In this work, we used a biologically realistic rate-based model to learn opposite neurons that exhibit experimentally observed tuning properties to bimodal stimuli and are topographically organized. Our learned neurons display contrast-invariant tuning, a widely observed tuning property of V1 neurons [11, 23], and their response to varying reliability of input stimulus agree with experimental observations qualitatively. Our model architecture is compatible with some existing decentralized models of multisensory integration, and therefore our work also provides a basis for learning such models in general.

Some studies of ventriloquism have explored the learning of the equivalent of congruent neurons in a similar decentralized model [9, 24, 25], but they assume *a priori* topographic organization of the multisensory neurons before learning. Here, no such assumption is made, and topographic organization naturally emerges via a Kohonen map-like mechanism [16]. They also did not explore the learning of opposite neurons, which is the key contribution of our study.

### Anti-Hebbian learning rule

Anti-Hebbian learning, or learning of the inhibitory connections, in our model differs from some other rate-based models in which anti-Hebbian learning is involved [12, 15, 32]. Instead of assigning a different learning rule to inhibitory neurons, our inhibitory neurons follow the same Hebbian learning rule as the excitatory neurons. We speculate that such a simple learning rule worked for us because of the delay we introduced to the inhibitory signal from congruent to opposite neurons, which is biologically realistic because of our model architecture. In fact, if such a delay is removed, opposite neurons cannot be learned well. We also tried using the anti-Hebbian learning rule introduced by Földiák, which takes the form Δ*w_ij_* = −*α*(*r_i_r_j_* − *p*^2^), for some fixed constant *p*. While opposite neurons can still be learned, the shape of the receptive fields can no longer be well-approximated by Gaussian or von-Mises distributions, an assumption in some decentralized models of multisensory integration [9, 24, 25, 29, 30].

### Causal Inference with opposite neurons

We are motivated by the theoretical observation that opposite neurons could provide a key step in causal inference by computing the Bayes factor [30, 31]. This is important because it is still unknown how the brain knows when to integrate or segregate multisensory cue information. However, the theoretical derivation assumes that opposite neurons simply sum up opposite inputs linearly, with no recurrent connections or divisive normalization among the opposite neurons. In contrast, our model is highly nonlinear [31]. Consequently, the theoretical derivations do not directly apply to the opposite neurons learned in our model. In future works, we aim to extend the theory to incorporate considerations of non-linearity in the circuit. Moreover, a possible future extension of this model would include a decision-making circuit that would determine when to integrate or segregation cue information based on the output of opposite neurons.

## Conclusion

We have demonstrated in this paper that our model can learn opposite neurons that generally agree with experimental observations. Our congruent and opposite neurons also learn a topographic organization via a Kohonen map-like mechanism. In addition, our model can be easily integrated with some existing multisensory integration models, paving the way towards a complete circuit for performing multisensory integration that can optimally decide whether to combine or segregate cue information.

## Materials and Methods

### Network Dynamics

The model assumes that each group of neurons (S1, S2, C1, C2, O1, O2) lies on a 1D ring, with each neuron’s position parameterized by *θ* ∈ [−*π*, *π*). The mean firing rate of neurons in the sensory input areas S1 and S2 is given by

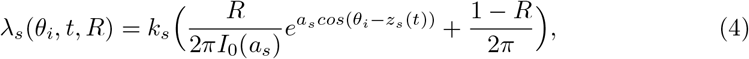

where *θ_i_* is the position of neuron *i* on the 1D ring. The subscript *s* ∈ {1, 2} indicates whether the input is from S1 or S2. *I*_0_(*x*) is the modified Bessel function of the first kind with order 0. *k_s_* is a scaling constant, while *R* ∈ [0,1] is the *reliability* of the input. For example, for a visual self-motion input, *R* = 0.5 would correspond to 50% reliability/coherence of the random dot stimulus. The reliability is kept to be 1 throughout the training. *a_s_* determines the width of the input, while *z_s_*(*t*) refers to the center of the input at time *t*.

This equation models the input as having the shape of a von-Mises distribution (first term of the equation) with a variable DC offset (second term of the equation). Von-Mises distribution can be thought of as an analogue of Gaussian distribution when the support is a circle. It is similar to the wrapped Gaussian distribution, which has been used to model the tuning of MSTd neurons [18]. Reliability controls the gain of the von-Mises distribution but does not affect the input width. This is consistent with the observation that tuning bandwidth of MT neurons (which provides visual input to MSTd neurons) is roughly “coherence-invariant,” meaning it is invariant to the reliability of the visual motion input [4]. The variable DC offset is set such that the total firing rate is independent of reliability, with 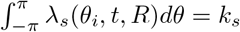. It is unclear whether MT neurons exhibit this property, but the requirement that total firing rate be relatively invariant to reliability is an important control of our model.

Let 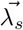 denote the vector of mean firing rate of neurons in S1 or S2. Let *W*_*c*1_, *W*_*c*2_, *W*_*o*1_, *W*_*o*2_, *W*_*o*1*c*1_, *W*_*o*2*c*2_ be the feedforward connections from S1 to C1, S2 to C2, S1 to O1, S2 to O2, C1 to O1 and C2 to O2, respectively. The direct feedforward input to congruent neurons is given by

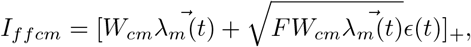

where *m* ∈ {1, 2} represents different modules, *ϵ*(*t*) is Gaussian noise with *μ* = 0, *σ* = 1, [*x*]_+_ = max(*x*, 0), and *F* = 1 is the Fano Factor. Similarly, the direct feedforward input to opposite neurons is given by

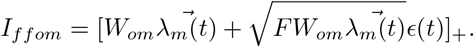

Recurrent connections among congruent and opposite neurons are modeled by

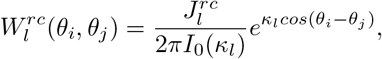

where *θ_i_* and *θ_j_* are positions of two different neurons *i* and *j* on the same ring. *l* ∈ {*c*1, *c*2, *o*1, *o*2} indicates whether the recurrent connection is among the ring of congruent neurons or opposite neurons. In theory, if we also model reciprocal connections in the same way as recurrent connections:

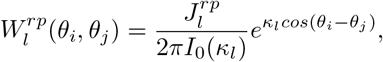

then 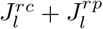 cannot be greater than a critical value *J_crit_*, or else the network can sustain spontaneous, consistent activities even after the removal of feedforward stimuli. The formula for *J_crit_* has been derived by Zhang et al. [30], and is given by

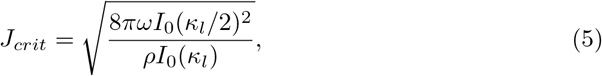

where *ρ* = *N*/2*π*. Although in our real implementations, the reciprocal connections are learned rather than set, we still imposed similar constraints to the upper bound of recurrent and reciprocal connections. We use *W_cm′,cm_* to denote the learned reciprocal connections from the congruent neurons in module *m* to those in module *m′*.

We did not explicitly model divisive normalization using neurons. Instead, the effects of divisive normalization among the ring *l* of neurons are directly incorporated into the calculation of firing rate:

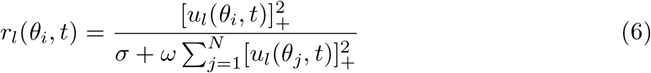

where *r_l_*(*θ_i_*) is the firing rate, *u_l_*(*θ_i_*) is the synaptic input, *N* is the number of neurons on ring *l, ω* is a constant that controls the strength of normalization, and *σ* adjusts the position of normalization. The normalization operation described here was used by Carandini and Heeger to model divisive normalization observed in biological data [6], and also in some previous studies of continuous attractor neural networks (CANN) [27, 28]. Experiments has also supported the presence of divisive normalization in multisensory integration areas [19]. We hypothesize that the operation could be carried out by a pool of inhibitory neurons. Again, we denote the vector of firing rates by 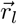 and the vector of synaptic inputs by 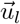.

Finally, the dynamics of congruent neurons and opposite neurons are given by

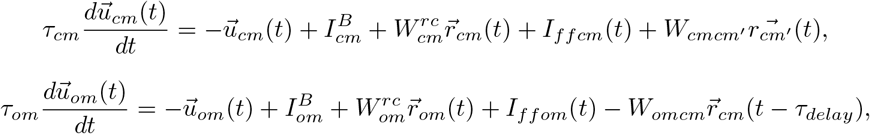

where *m* ∈ {1, 2} indicates specific modules. The first term for both equations is a decay term. The second term 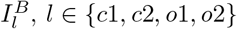 is a constant background input. The third term corresponds to input from recurrent connections within the ring of congruent neurons or opposite neurons, and 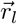 is given by Eq. 6. The fourth term correspond to direct feedforward inputs from uni-sensory neurons.

The last term for congruent neurons represent the reciprocal input from the other module. The last term for opposite neurons has a negative sign in front of 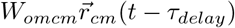 since the connection is inhibitory as well as a delay *τ_delay_* of the signal from congruent neurons to opposite neurons. This delay is essential for the learning of opposite neurons, for otherwise the excitatory and inhibitory input will keep canceling out throughout training, which in turn degrades the learning efficacy of opposite neurons. We note that this delay is also biologically plausible, since the inhibition, which may go through interneurons, is disynaptic, and would therefore need longer time to transmit in comparison to the monosynaptic excitatory input.

After each update, we rectify the weights with [*w_ij_*]_+_ to ensure all weights are non-negative.

### Learning Rules

The network learns the feedforward excitatory and inhibitory weights via the same local, Hebbian learning rule, with

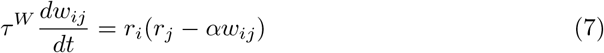

where *w_ij_* denote an excitatory/inhibitory connection from neuron *j* to neuron *i*. As mentioned, we also enforce the constraint that all weights must be non-negative. Note that this is *not* Oja’s rule, where the second term inside the bracket would be *r_i_w_ij_*. Ursino et al. showed that for a simplified model without recurrent connections, this learning rule (with *α* = 1) for excitatory neurons allows the receptive field of the neuron to match its average input, which in turn allows maximum likelihood estimation in multisensory integration to be performed simply by reading out the position of the neuron with maximal firing rate [25].

### Simulation Details

There were *N* = 180 neurons on each of the six rings of neurons (S1, S2, C1, C2, O1, O2), distributed uniformly over the stimulus space [−*π*, *π*). We set the synaptic input time constants to be *τ_c_* = *τ_o_* = *τ* = 10. Although this number is unitless, one can relate it to 10 millisecond (ms). Euler’s method was used with a step size of Δ_*t*_ = 0.1*τ*, and the simulation was run for *T* = 180000 = 180000Δ_*t*_. The learning rule time constant was 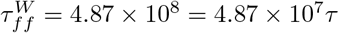 for feedforward weights (direct and indirect), and *α_ff_* = 9740. Since reciprocal weights are subject to constraints in Eq. 5, they are in general smaller than feedforward weights, hence we set 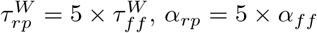. Here we note that the ratio between firing rate and weight was chosen empirically, other ratios may also give rise to qualitatively similar simulated results with the related parameters re-tuned.

For sensory inputs, *a*_1_ = *a*_2_ = 1.5, and *k*_1_ = *k*_2_ = 2*πI*_0_(3)*e*^−3^ ≈ 1.5. The position of input from S1, *z*_1_(*t*), was generated by first randomly permuting an evenly spaced sequence of inputs from −*π* to *π*, then adding a Gaussian noise with *μ* = 0 and *σ* = 2°, each lasting *τ_stim_* = 100 = 10*τ*. *z*_2_(*t*) was generated by adding another Gaussian noise with the same *μ* and *σ* to *z*_1_(*t*). The Fano Factor *F* of the summed feedforward input was set to 1. For recurrent connections, *κ*_*c*1_ = *κ*_*c*2_ = *κ*_*o*1_ = *κ*_*o*2_ = 3; for divisive normalization, *ω* = 2.46 ·10^-4^, *σ* = 0.3.

All synaptic inputs were initialized to 0. The feedforward weights were all initialized with the following method: Consider feedforward connections from a pool of input neurons indexed by *j* to a pool of target neurons indexed by *i*. For each *j*, we sample 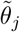 from a uniform distribution over the *N* target neurons without replacement (i.e. 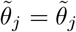 if and only if *j* = *j′*), as well as a multiplicative factor *A_j_* from a log normal distribution with arithmetic mean of 0.3 and arithmetic variance of 0.1. Then the initial feedforward connections are given by

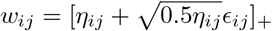

where

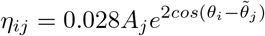

and *ϵ_ij_* is i.i.d. Gaussian noise with *μ* = 0 and *σ* = 1. Intuitively, this models each input neuron as projecting to a random target location with variable connection strength and a spatial spread given by von Mises distribution. The initial reciprocal connections are set to be 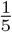 times of feedforward connections.

### Calculating the concentration of peak distribution

Let *N_t_* denote the number of simulations when deriving the peak distribution. The concentration of the distribution is calculated to be

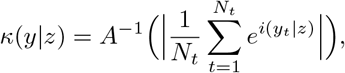

where *A* is given by

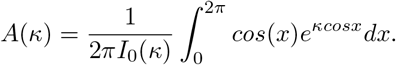

## References

1. Angelaki DE, Gu Y, DeAngelis GC. Neural correlates of multisensory cue integration in macaque MSTd. Nature Neuroscience. 2008;11(10):1201–1210. doi:10.1038/nn.2191.

2. Beck JM, Pouget A, Latham PE, Ma WJ. Bayesian inference with probabilistic population codes. Nature Neuroscience. 2006;9(11):1432–1438. doi:10.1038/nn1790.

3. Bertin R, Berthoz A. Visuo-vestibular interaction in the reconstruction of travelled trajectories. Experimental Brain Research. 2004;154(1):11–21. doi:10.1007/s00221-003-1524-3.

4. Britten KH, Newsome WT. Tuning Bandwidths for Near-Threshold Stimuli in Area MT. Journal of Neurophysiology. 1998;80(2):762–770. doi:10.1152/jn.1998.80.2.762.

5. Britten KH, Shadlen MN, Newsome WT, Movshon JA. Responses of neurons in macaque MT to stochastic motion signals. Visual Neuroscience. 1993;10(6):1157–1169. doi:10.1017/S0952523800010269.

6. Carandini M, Heeger DJ. Normalization as a canonical neural computation. Nature reviews Neuroscience. 2011;13(1):51–62. doi:10.1038/nrn3136.

7. Chen A, DeAngelis GC, Angelaki DE. Representation of vestibular and visual cues to self-motion in ventral intraparietal cortex. Journal of Neuroscience. 2011; 31(33): 12036–12052.

8. Chen A, Deangelis GC, Angelaki DE. Functional Specializations of the Ventral Intraparietal Area for Multisensory Heading Discrimination. The Journal of neuroscience: the official journal of the Society for Neuroscience. 2013;33(8):3567–3581. doi:10.1523/JNEUROSCI.4522-12.2013.

9. Cuppini C, Shams L, Magosso E, Ursino M. A biologically inspired neurocomputational model for audiovisual integration and causal inference. European Journal of Neuroscience. 2017;46(9):2481–2498. doi:10.1111/ejn.13725.

10. Dokka K, Park H, Jansen M, DeAngelis GC, Angelaki DE. Causal inference accounts for heading perception in the presence of object motion. Proceedings of the National Academy of Sciences of the United States of America. 2019;116(18):9060–9065. doi:10.1073/pnas.1820373116.

11. Finn IM, Priebe NJ, Ferster D. The Emergence of Contrast-Invariant Orientation Tuning in Simple Cells of Cat Visual Cortex. Neuron. 2007;54(1):137–152. doi:10.1016/j.neuron.2007.02.029.

12. Földiák P. Forming sparse representations by local anti-Hebbian learning. Biological cybernetics. 1990;64(2):165–170. doi:10.1007/BF00202929.

13. Gu Y, Deangelis GC, Angelaki DE. Causal Links between Dorsal Medial Superior Temporal Area Neurons and Multisensory Heading Perception. The Journal of neuroscience: the official journal of the Society for Neuroscience. 2012;32(7):2299–2313. doi:10.1523/JNEUROSCI.5154-11.2012.

14. Gu Y, Watkins PV, Angelaki DE, DeAngelis GC. Visual and Nonvisual Contributions to Three-Dimensional Heading Selectivity in the Medial Superior Temporal Area. Journal of Neuroscience. 2006;26(1):73–85. doi:10.1523/JNEUROSCI.2356-05.2006.

15. King PD, Zylberberg J, DeWeese MR. Inhibitory Interneurons Decorrelate Excitatory Cells to Drive Sparse Code Formation in a Spiking Model of V1. The Journal of neuroscience: the official journal of the Society for Neuroscience. 2013;33(13):5475–5485. doi:10.1523/JNEUROSCI.4188-12.2013.

16. Kohonen T. Self-organized formation of topologically correct feature maps. Biological Cybernetics. 1982;43(1):59–69. doi:10.1007/BF00337288.

17. Körding KP, Beierholm U, Ma WJ, Quartz S, Tenenbaum JB, Shams L. Causal Inference in Multisensory Perception. PLoS One. 2007;2(9):e943. doi:10.1371/journal.pone.0000943.

18. Morgan ML, DeAngelis GC, Angelaki DE. Multisensory Integration in Macaque Visual Cortex Depends on Cue Reliability. Neuron. 2008;59(4):662–673. doi:10.1016/j.neuron.2008.06.024.

19. Ohshiro T, Angelaki DE, DeAngelis GC. A Neural Signature of Divisive Normalization at the Level of Multisensory Integration in Primate Cortex. Neuron. 2017;95(2):39–411.e8. doi:10.1016/j.neuron.2017.06.043.

20. Pearl J. Causal inference in statistics: An overview. Statistics Surveys. 2009;3:96–146. doi:10.1214/09-SS057.

21. Sato Y, Toyoizumi T, Aihara K. Bayesian Inference Explains Perception of Unity and Ventriloquism Aftereffect: Identification of Common Sources of Audiovisual Stimuli. Neural Computation. 2007;19(12):3335–3355. doi:10.1162/neco.2007.19.12.3335.

22. Shams L, Beierholm UR. Causal inference in perception. Trends in Cognitive Sciences. 2010;14(9):425–432. doi:10.1016/j.tics.2010.07.001.

23. Troyer TW, Krukowski AE, Priebe NJ, Miller KD. Contrast-Invariant Orientation Tuning in Cat Visual Cortex: Thalamocortical Input Tuning and Correlation-Based Intracortical Connectivity. Journal of Neuroscience. 1998;18(15):5908–5927. doi:10.1523/JNEUROSCI.18-15-05908.1998.

24. Ursino M, Crisafulli A, di Pellegrino G, Magosso E, Cuppini C. Development of a Bayesian Estimator for Audio-Visual Integration: A Neurocomputational Study. Frontiers in computational neuroscience. 2017;11:89. doi:10.3389/fncom.2017.00089.

25. Ursino M, Cuppini C, Magosso E. Multisensory Bayesian Inference Depends on Synapse Maturation during Training: Theoretical Analysis and Neural Modeling Implementation. Neural computation. 2017;29(3):735–782.

26. Wallace MT, Roberson GE, Hairston WD, Stein BE, Vaughan JW, Schirillo JA. Unifying multisensory signals across time and space. Experimental Brain Research. 2004;158(2):252–258. doi:10.1007/s00221-004-1899-9.

27. Wu S, Hamaguchi K, ichi Amari S. Dynamics and Computation of Continuous Attractors. Neural Computation. 2008;20(4):994–1025. doi:10.1162/neco.2008.10-06-378.

28. Wu S, Wong KYM, Fung CCA, Mi Y, Zhang W. Continuous Attractor Neural Networks: Candidate of a Canonical Model for Neural Information Representation [version 1; referees: 2 approved]. F1000Research. 2016;5:156. doi:10.12688/f1000research.7387.1.

29. Zhang WH, Chen A, Rasch MJ, Wu S. Decentralized Multisensory Information Integration in Neural Systems. The Journal of neuroscience: the official journal of the Society for Neuroscience. 2016;36(2):532–547. doi:10.1523/JNEUROSCI.0578-15.2016.

30. Zhang WH, Wang H, Chen A, Gu Y, Lee TS, Wong KM, et al. Complementary congruent and opposite neurons achieve concurrent multisensory integration and segregation. eLife. 2019;8. doi:10.7554/eLife.43753.

31. Zhang WH, Wu S, Doiron B, Lee TS. A Normative Theory for Causal Inference and Bayes Factor Computation in Neural Circuits;.

32. Zylberberg J, Murphy JT, DeWeese MR. A Sparse Coding Model with Synaptically Local Plasticity and Spiking Neurons Can Account for the Diverse Shapes of V1 Simple Cell Receptive Fields. PLoS computational biology. 2011;7(10):e1002250. doi:10.1371/journal.pcbi.1002250.

